# Scalable probabilistic PCA for large-scale genetic variation data

**DOI:** 10.1101/729202

**Authors:** Aman Agrawal, Alec M. Chiu, Minh Le, Eran Halperin, Sriram Sankararaman

## Abstract

Principal component analysis (PCA) is a key tool for understanding population structure and controlling for population stratification in genome-wide association studies (GWAS). With the advent of large-scale datasets of genetic variation, there is a need for methods that can compute principal components (PCs) with scalable computational and memory requirements. We present ProPCA, a highly scalable method based on a probabilistic generative model, which computes the top PCs on genetic variation data efficiently. We applied ProPCA to compute the top five PCs on genotype data from the UK Biobank, consisting of 488,363 individuals and 146,671 SNPs, in less than thirty minutes. Leveraging the population structure inferred by ProPCA within the White British individuals in the UK Biobank, we scanned for SNPs that are not well-explained by the PCs to identify several novel genome-wide signals of recent putative selection including missense mutations in RPGRIP1L and TLR4.

**Author Summary:** Principal component analysis is a commonly used technique for understanding population structure and genetic variation. With the advent of large-scale datasets that contain the genetic information of hundreds of thousands of individuals, there is a need for methods that can compute principal components (PCs) with scalable computational and memory requirements. In this study, we present ProPCA, a highly scalable statistical method to compute genetic PCs efficiently. We systematically evaluate the accuracy and robustness of our method on large-scale simulated data and apply it to the UK Biobank. Leveraging the population structure inferred by ProPCA within the White British individuals in the UK Biobank, we identify several novel signals of putative recent selection.

## Introduction

Inference of population structure is a key step in population genetic analyses [1] with applications that include understanding genetic ancestry [2–4] and controlling for confounding in genome-wide association studies (GWAS) [5]. While several methods have been proposed to infer population structure (e.g., [6–10]), principal component analysis (PCA) is one of the most widely used [11, 6]. Unfortunately, the naive approach for estimating principal components (PCs) by computing a full singular value decomposition (SVD) scales quadratically with sample size (for datasets where the number of SNPs is larger than sample size), resulting in runtimes unsuitable for large data sets.

In light of these challenges, several solutions have been proposed for the efficient computation of PCs. One approach taken by two recent scalable implementations (FastPCA [12] and Flash-PCA2 [13]) takes advantage of the fact that typical applications of PCA in genetics only require computing a small number of top PCs; *e.g.* GWAS typically use 5-20 PCs to correct for stratification [14]. An alternative approach for efficient computation of PCs takes advantage of the parallel computation infrastructure of the cloud [15]. However, the cost of cloud usage is roughly proportional to the number of CPU hours used by these algorithms, making them cost-prohibitive. Finally, these scalable implementations lack a full probabilistic model, making them challenging to extend to settings with missing genotypes or linkage disequilibrium (LD) between SNPs.

In this work, we describe ProPCA, a scalable method to compute the top PCs on genotype data. ProPCA is based on a previously proposed probabilistic model [16, 17], of which PCA is a special case. While PCA treats the PCs and the PC scores as fixed parameters, probabilistic PCA imposes a prior on the PC scores. This formulation leads to an iterative Expectation Maximization (EM) algorithm for computing the PCs. ProPCA leverages the structure of genotype data so that each iteration of the EM algorithm can be computed in time that scales sub-linear in the number of individuals or SNPs. The EM algorithm requires only a small number of iterations to obtain accurate estimates of the PCs resulting in a highly scalable algorithm.

In both simulated and real data, ProPCA is able to accurately infer the top PCs while scaling favorably with increasing sample size. We applied ProPCA to compute the top five PCs on genotype data from the UK Biobank, consisting of 488,363 individuals and 146,671 SNPs, in less than thirty minutes. Leveraging the population structure inferred by ProPCA within the White British individuals in the UK Biobank [18], we scanned for SNPs that are not well-modeled by the top PCs to identify several novel genome-wide signals of recent positive selection. Our scan recovers sixteen loci that are highly differentiated across the top five PCs that are likely signals of recent selection. While these loci include previously reported targets of selection [12], the larger sample size that we analyze here allows us to identify eleven novel signals including a missense mutation in RPGRIP1L (*p* = 2.09 × 10^*-*9^) and another in TLR4 (*p* = 7.60 × 10^*-*12^).

A number of algorithms that analyze genotype data, including methods for heritability estimation and association testing, can be modeled as iterative procedures where the core computational operation is similar to that solved by ProPCA. Thus, the algorithm that we employ in this work can potentially lead to highly scalable algorithms for a broad set of population genetic analyses.

## Results

### Accuracy

We first assessed the accuracy of ProPCA using the simulation framework described in the Methods. We generated datasets containing 50, 000 SNPs and 10, 000 individuals across *q* populations, where *q* was chosen to be 5 and 10. The populations were simulated with varying levels of population differentiation that are typical of present-day human populations (values of *F*_*st*_ ranging from 0.001 to 0.01) and were small enough so that we could compute the full SVD thereby allowing us to estimate the accuracy of the PCs computed by ProPCA. To measure accuracy, we computed the mean of explained variances (MEV), a measure of the overlap between the subspaces spanned by the PCs estimated by ProPCA compared to the PCs computed using a full SVD (Methods). ProPCA estimates highly accurate PCs (values of MEV close to 1) across the range of parameters (Table 1).

**Table 1:** ProPCA accurately estimates principal components: The principal components computed by ProPCA are compared to the PCs obtained from a full SVD on a genotype dataset containing 50, 000 SNPs and 10, 000 individuals. Accuracy was measured by the mean of explained variance (MEV) which measures the overlap between the set of PCs inferred from ProPCA and those from SVD across values of *F*_*st*_ ∈ {0.001, …, 0.01}. We report MEV for *K* = 5 using 5 populations as well as for *K* = 10 PCs using 10 populations.

| $F_{st}$ | MEV | |
| --- | --- | --- |
| | $K = 5$ | $K = 10$ |
| 0.001 | 0.987 | 1.000 |
| 0.002 | 0.999 | 1.000 |
| 0.003 | 0.999 | 1.000 |
| 0.004 | 0.999 | 1.000 |
| 0.005 | 1.000 | 1.000 |
| 0.006 | 1.000 | 1.000 |
| 0.007 | 1.000 | 1.000 |
| 0.008 | 1.000 | 1.000 |
| 0.009 | 1.000 | 1.000 |
| 0.010 | 1.000 | 1.000 |

### Runtime

We assessed the scalability of ProPCA with increasing sample size (Methods). We simulated genotypes from five populations containing 100, 000 SNPs and sample sizes varying from 10, 000 to 1, 000, 000 with *F*_*st*_ = 0.10.

We compared the wall-clock time for running ProPCA, the SVD implementation in PLINK (PLINK SVD [19]), FastPCA [12] and FlashPCA2 [13]. The SVD implementation in PLINK could not run in reasonable time on datasets exceeding 70, 000 individuals (Figure 1a). While FastPCA, FlashPCA2 and ProPCA all scale with sample size, ProPCA is about four times faster than FastPCA and twice as fast as FlashPCA2 (Figure 1b). ProPCA computes PCs in about 30 minutes even on the largest data containing a million individuals and 100, 000 SNPs.

**Figure 1:**
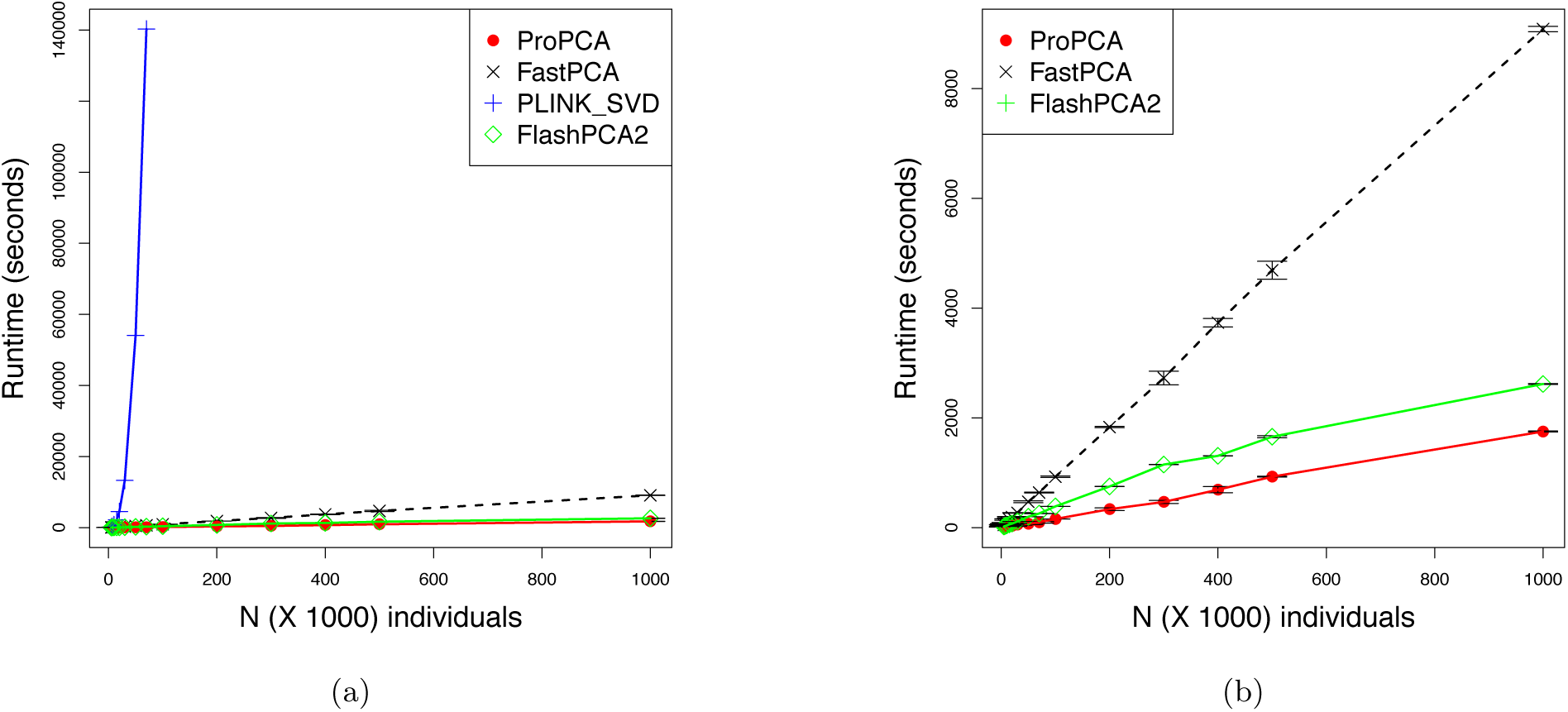
ProPCA is computationally efficient: Comparison of runtimes over simulated genotype data containing 100, 000 SNPs, six subpopulations, *F*_*st*_ = 0.10 and individuals varying from 10, 000 to 1, 000, 00. We report the mean and standard deviation over ten trials. Figures 1a and 1b display the total runtime (capped to a maximum of 100 hours and a maximum memory of 64 GB). Figure 1b compares the runtimes of all algorithms excluding PLINK SVD which could only run successfully up to a sample size of 70, 000.

Since ProPCA, FastPCA, and FlashPCA2 are all based on iterative algorithms, their runtimes depend on details of convergence criterion. We performed an additional experiment to compare the runtime of ProPCA and FastPCA (for which we could instrument the source code) for a single iteration and found ProPCA to be three to four times faster than FastPCA across the range of sample sizes (Figure S1). Measuring the accuracy of the PCs (MEV) as a function of runtime (on datasets with a range of *F*_*st*_ containing 50, 000 SNPs and 10, 000 individuals so that we could compare the estimated PCs to exact PCs), ProPCA attains a given MEV in about half the time as FastPCA (Figure 2). We were unable to include FlashPCA2 in these comparisons as we were unable to instrument the source code to control the number of iterations.

**Figure 2:**
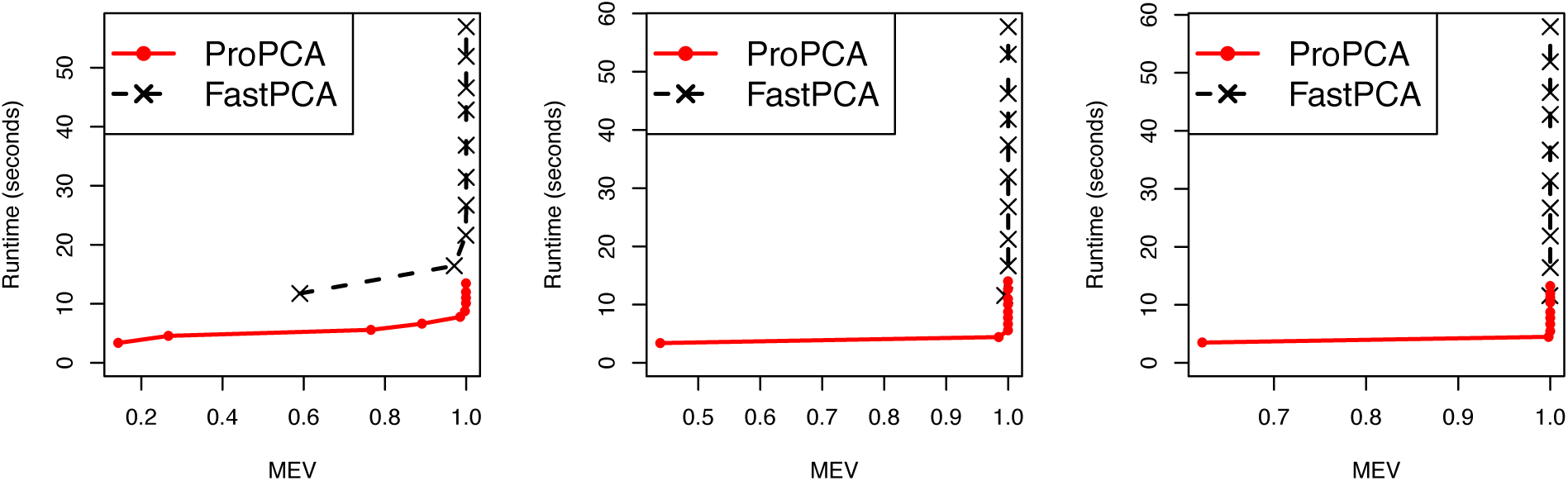
ProPCA is computationally efficient relative to FastPCA: We compute the total time taken to estimate the top five principal components as a function of a measure of accuracy (MEV) for ProPCA compared to FastPCA. We performed these comparisons on simulated genotype data containing 50, 000 SNPs, 10, 000 individuals, six subpopulations, and *F*_*st*_ ∈ {0.001, 0.005, 0.10}. We were unable to leverage the source code of FlashPCA2 to include in these comparisons.

### Application to real genotype data

We applied ProPCA to genotype data from Phase 1 of the 1000 Genomes project [20]. On a dataset of 1092 individuals and 442, 350 SNPs, ProPCA computes the top five PCs with an MEV of 0.968 producing PCs that are qualitatively indistinguishable from running a full SVD (Figure S4, Table S1). We also applied ProPCA to genotype data from the UK Biobank [18] consisting of 488, 363 individuals and 146, 671 SNPs after QC. ProPCA can compute the top five PCs in about 27 minutes and the resulting PCs reflect population structure within the UK Biobank, consistent with previous studies [18] (Figure 3a).

**Figure 3:**
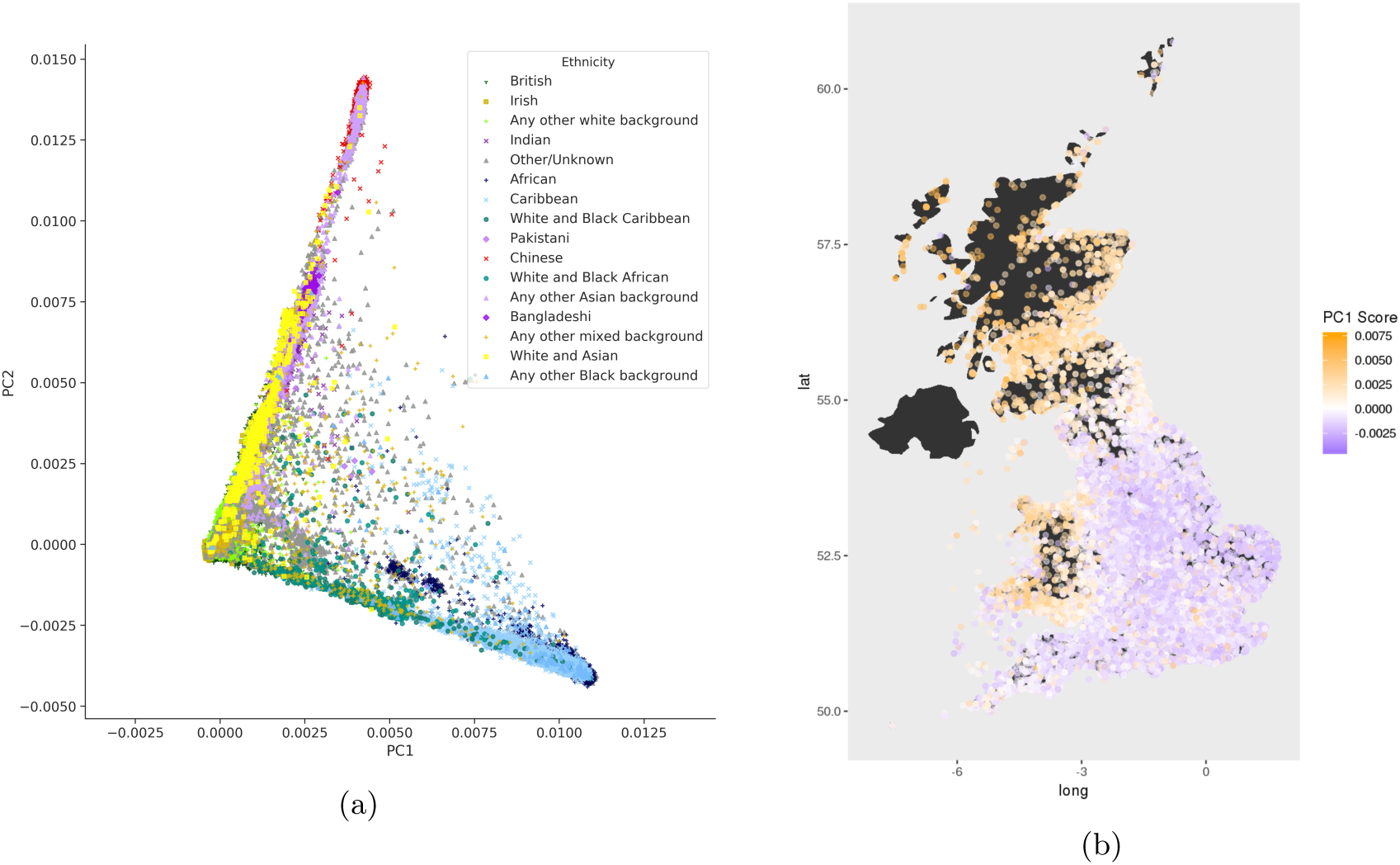
Principal components uncover population and geographic structure in the UK Biobank: We used ProPCA to compute PCs on the UK Biobank data. Figure 3a shows the first two principal components to reveal population structure. Figure 3b shows geographic structure by plotting the score of 276, 736 unrelated White British individuals on the first principal component on their birth location coordinates.

### Application to scans for selection

We developed a statistical test to search for SNPs that are not well-modeled by the ProPCA model as a means of discovering signals of natural selection (Methods). This statistic relies on the observation that a SNP evolving under positive selection is expected to exhibit differentiation in the frequencies of its alleles that is extreme compared to a typical SNP that is evolving neutrally [21]. Since deviations from the ProPCA model can occur due to reasons unrelated to selection, we filtered out SNPs with high rates of missingness, low minor allele frequency (MAF), and presence in regions of long-range LD [22](Methods). We ran ProPCA to infer the top five PCs on 276, 736 unrelated White British samples and the UK Biobank SNP set consisting of 146, 671 SNPs obtained by further removing SNPs in high LD (Figure S5).

The Pearson correlation coefficient between birth location coordinates and the PC score for each individual reveals that the estimated PCs capture geographic structure within the UK (Figure 3b, Figure S6, Table S2). We used these PCs to perform a selection scan on a larger set of 516, 140 SNPs and we report SNPs that are genome-wide significant after accounting for the number of SNPs as well as PCs tested (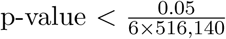; we use 6 to account for the additional combined test statistic that we describe later). We ensured that the selection statistic for each PC was well-calibrated against a 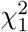 distribution (Fig. S7) and genomic inflation (*λ*_*GC*_) values for each of the PCs showed no substantial inflation (Table S3). While our statistic is closely related to a previously proposed statistic to detect selection on PCs [12], we found that our proposed statistic is better calibrated (Table S3).

Our scan revealed a total of 59 SNPs that were genome-wide significant (Table S4). Clustering these signals into 1 Mb windows centered around the most significant SNP for each PC, we obtained twelve non-overlapping loci that contain putative targets of selection (Figure 4, Table S5). These twelve loci include five that were previously reported to be signals of selection in the UK with genome-wide significance: *LCT* (rs7570971 with *p* = 8.51 × 10^*-*16^), *TLR1* (rs5743614, *p* = 5.65 × 10^*-*25^), *IRF4* (rs62389423, *p* = 8.80 × 10^*-*42^), *HLA* (rs9267817, *p* = ×6.17 × 10^*-*9^), and *FUT2* (rs492602, *p* = 7.02 × 10^*-*10^) [12]. The larger sample size that we analyze here also reveals novel signals at additional loci. Four of the twelve signals were previously suggested to be signals of selection but were not genome-wide significant: *HERC2* (rs12913832, *p* = 5.21×10^*-*10^), *RPGRIP1L* (rs61747071, *p* = 2.09 × 10^*-*9^), *SKI* (rs79907870, *p* = 2.58 × 10^*-*9^), *rs77635680* (*p* = 2.22 × 10^*-*10^) [12] while the remaining three loci: *HERC6* (rs112873858, *p* = 2.68 × 10^*-*11^), *rs6670894* (*p* = 4.98 × 10^*-*9^), and *rs12380860* (*p* = 8.62 × 10^*-*9^) appear to be previously unreported (Section S7).

**Figure 4:**
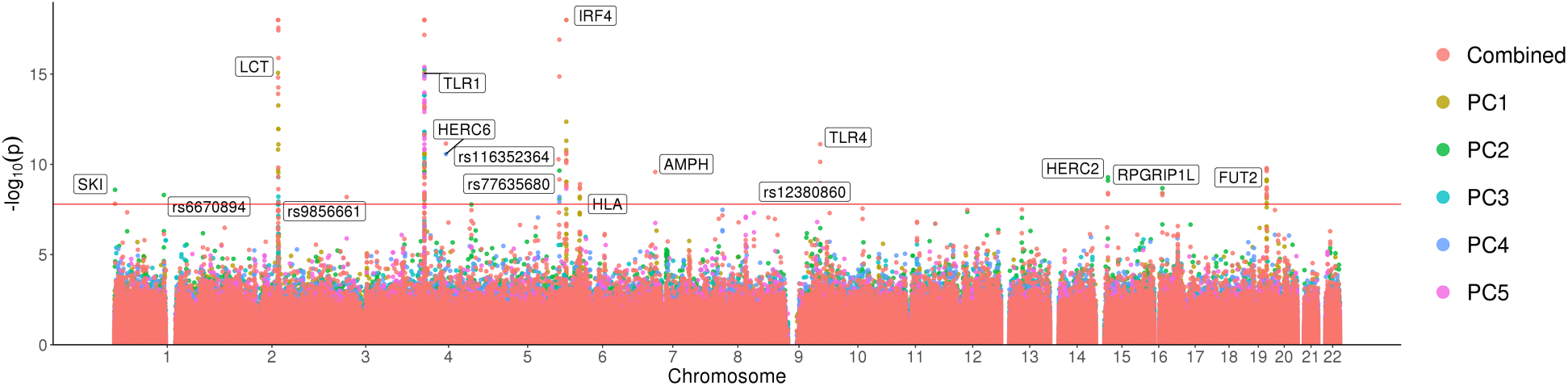
Selection scan for the first five principal components in the white British individuals in the UK Biobank: A Manhattan plot with the − log_10_ *p* values associated with the test of selection displayed for the first five principal components for the unrelated White British subset of the UK Biobank. The red line represents the Bonferroni adjusted significance level (*α* = 0.05). Significant loci are labeled. Signals above − *log*_10_(*p*) = 18 were capped at this value for better visualization.

To validate our findings, we utilized birth location coordinates for each individual and assigned them to geographical regions in the UK as defined in the Nomenclature of Territorial Units for Statistics level 3 (NUTS3) classification. We performed a test of association between the allele frequency of the top SNP in each of our novel loci with geographical regions (Table S9) and confirmed that SNPs identified in our selection scan show differences in allele frequencies across specific geographical regions (Table S9).

One of the novel genome-wide significant loci is *RPGRIP1L. RPGRIP1L* is a highly conserved gene that encodes a protein that localizes at primary cilia and is important in development [23]. Mutations in this gene have been implicated with neurological disorders such as Joubert syndrome and Meckel syndrome [24], conditions that sometimes also result in additional symptoms such as eye diseases and kidney disorders [25]. The SNP with the most significant p-value in our scan in *RPGRIP1L*, rs61747071, is a missense loss-of-function mutation A229T that has been shown to lead to photoreceptor loss in ciliopathies [26].

We created an additional variant of our selection statistic which tests for SNPs that are not well-modeled by a linear combination of the first five PCs by summing the per-PC 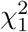 statistics resulting in a new chi-squared statistic with five degrees of freedom. Combining signals across PCs has been previously shown to boost power in association testing [27]. We verified that the resulting combined statistic is also calibrated (Fig S7 and Table S3). Under this combined statistic, we recover majority of the loci found on each individual PC, but we also discover four additional novel loci: *AMPH* (rs118079376, *p* = 2.64 × 10^*-*10^), *TLR4* (rs4986790, *p* = 7.60 × 10^*-*12^), *rs9856661* (*p* = 6.46 × 10^*-*9^), and *rs116352364* (*p* = 5.24 × 10^*-*11^) (Table S8).

*TLR4* is a member of the toll-like receptor family. The TLR gene family is known to play a fundamental role in pathogen recognition and activation of innate immunity, but TLR4 in particular is involved with proinflammatory cytokines and has a pro-carcinogenic function [28]. The SNP with the most significant *p*-value at our *TLR*4 locus is rs4986790, a missense D299G mutation and D259G mutation on two different transcripts for the TLR4 gene. The D299G mutation is of particular interest as this mutation is strongly correlated with increased infection by *Plasmodium falciparum*, a parasite that causes malaria [29, 30]. Additional details on the signals of selection can be found in the Supplementary Information (SI Section S7).

## Discussion

We have presented, ProPCA, a scalable method for PCA on genotype data that relies on performing inference in a probabilistic model. Inference in this model consists of an iterative procedure that uses a fast matrix-vector multiplication algorithm. We have demonstrated its accuracy and efficiency across diverse settings. Further, we have demonstrated that ProPCA can accurately estimate population structure within the UK Biobank dataset and how this structure can be leveraged to identify targets of recent putative selection.

The algorithm that we employ here to accelerate the EM updates is of independent interest. Beyond PCA, several algorithms that operate on genotype data perform repeated matrix-vector multiplication on the matrix of genotypes. For example, association tests and permutation tests, can be formulated as computing a matrix-vector product where the matrix is the genotype matrix while the vector consists of phenotype measurements. The idea that SVD computations can leverage fast matrix-vector multiplication operations to obtain computational efficiency is well known in the numerical linear algebra literature [31]. Indeed, the algorithms [31] implemented in FastPCA [12] as well as FlashPCA2 [13] can also utilize these ideas to gain additional computational efficiency. Alternate approaches to improve matrix-vector multiplication in the genetics setting include approaches that rely on sparsity of the genotype matrix. It is important to note that the speedup obtained from the Mailman algorithm does not rely explcitly on sparsity and could be applied even to dense matrices. It would be of interest to contrast these approaches and to investigate the potential to combine to leverage sparsity as well as the discrete nature of the genotype matrix.

The probabilistic formulation underlying ProPCA allows the algorithm to be generalized in several directions. One direction is the application of PCA in the presence of missing data that often arises when analyzing multiple datasets. We have explored an extension of the ProPCA model to this setting (SI Section S5). While this approach is promising, a limitation of the use of the Mailman algorithm within ProPCA is the requirement of discrete genotypes, which prevents ProPCA from being directly applied to dosages. Another potential future direction is in modeling linkage disequilibrium and in incorporating rare variants which have the potential to reveal structure that is not apparent from the analysis of common SNPs [32, 33]. Current applications of PCA remove correlated SNPs and singletons though this has been shown to discard information [12]. One possible way to incorporate LD would leverage the connection between haplotype copying models [34] and the multivariate normal model of PCA [35], or by a whitening transformation [4]. Further, the observation model can also be modified to account for the discrete nature of genotypes [3, 36]. A number of non-linear dimensionality reduction methods have been recently proposed [37, 38]. A comparison of these methods to ProPCA (in terms of statistical structure that the methods aim to detect, robustness and ability to handle missing data, as well computational scalability) would be of great interest. Finally, leveraging fine-scale population structure inferred from large-scale data to study recent positive selection in human history is an important direction for future work. The challenge is to design realistic statistical models of population structure while enabling inference at scale.

ProPCA is available at https://github.com/sriramlab/ProPCA.

## Material and Methods

### Principal Components Analysis (PCA)

We observe genotypes from *n* individuals at *m* SNPs. The genotype vector for individual *i* is a length *m* vector denoted by ***g***_*i*_ ∈ {0, 1, 2}^*m*^. The *j*^*th*^ entry of ***g***_*i*_ denotes the number of minor allele carried by individual *i* at SNP *j*. Let ***G*** be the *m* × *n* genotype matrix where ***G*** = [***g***_1_ … ***g***_*n*_]. Let ***Y*** denote the matrix of standardized genotypes obtained by centering and rescaling each row of the genotype matrix ***G*** so that 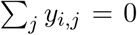 and 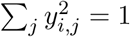 for all *i* ∈ {1, …, *m*}.

Principal components analysis (PCA) [11] attempts to find a low-dimensional linear transformation of the data that maximizes the projected variance or, equivalently, minimizes the reconstruction error. Given the *m* × *n* matrix ***Y*** of standardized genotypes and a target dimension *k*, PCA attempts to find a *m* × *k* matrix with orthonormal columns ***W*** and *n* × *k* matrix ***Z*** that minimizes the reconstruction error: ‖***Y*** *-* ***WZ***^T^‖_*F*_ where 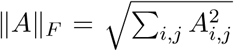 is the Frobenius norm of the matrix ***A***. To solve the PCA problem, we perform a singular-value decomposition (SVD) of the standardized genotype matrix ***Y*** = ***UΣV***^T^ and set 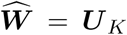, where ***U***_*K*_ is a *m* × *k* matrix containing the *k* columns of ***U*** corresponding to the *k* largest singular vectors of ***Y***.

### Probabilistic PCA

PCA can be viewed as a limiting case of the probabilistic PCA model [10, 16, 17]. Probabilistic PCA models the observed data ***y***_*i*_ ∈ ℝ^*m*^,*i* ∈ {1 …, *n*} as a linear transformation of a *k*-dimensional latent random variable ***x***_*i*_ (*k* ≤ *m*) with additive Gaussian noise. Denoting the linear transformation by the *m × k* matrix ***C***, and the (*m*-dimensional) noise by ***ϵ***_*i*_ (with isotropic covariance matrix *σ*^2^***I***_*m*_), the generative model can be written as

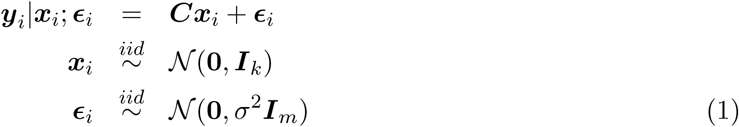

The maximum likelihood estimate of the matrix ***C*** in this model has been shown to span the *k*-dimensional principal subspace of the data ***Y*** = [***y***_1_, … ***y***_*n*_] [39].

### EM algorithm for PCA

Since probabilistic PCA is a probabilistic model endowed with latent variables, the EM algorithm presents a natural approach to compute the maximum likelihood estimates of the model parameters (***C***, *σ*^2^) [16, 17]. The EM algorithm for learning the principal components can be derived as a special case of the EM algorithm for the probabilistic PCA model where the variance of the observation noise *σ*^2^ tends to zero leading to these updates:

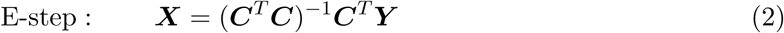

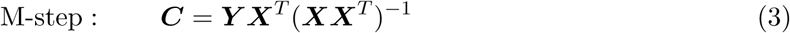

Here ***X*** = [***x***_1_ … ***x***_*n*_] is a *k* × *n* matrix and ***Y*** = [***y***_1_ … ***y***_*n*_] is a *m* × *n* matrix. Noting that all matrix inversions require inverting a *k* × *k* matrix, the computational complexity of the E-step is 𝒪(*k*^2^*m* + *k*^3^ + *k*^2^*m* + *mnk*) while the computational complexity of the M-step is 𝒪(*k*^2^*n* + *k*^3^ + *k*^2^*n* + *mnk*). For small *k* and large *m, n*, the per-iteration runtime complexity is 𝒪(*mnk*). Thus, the EM algorithm provides a computationally efficient estimator of the top *k* PCs when the number of PCs to be estimated is small.

### Sub-linear time EM

The key bottleneck in the EM algorithm is the multiplication of the matrix ***Y*** with matrices ***E*** = (***C***^*T*^ ***C***)^*-*1^***C***^*T*^ and ***M*** = ***X***^*T*^ (***XX***^*T*^)^*-*1^.

The vectors representing the sample mean and standard deviation of the genotypes at each SNP are denoted 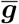 and ***s***. Assuming no entry in ***s*** is zero (we remove SNPs that have no variation across samples), the matrix of standardized genotypes ***Y*** can be written as:

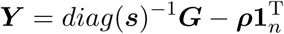

Here *diag*(***x***) is an operator that constructs a diagonal matrix with the entries of ***x*** along its diagonals, 1_*n*_ is a length *n* vector with each entry equal to one, and ***ρ*** is a length *m* vector with 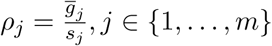

The EM updates can be written as:

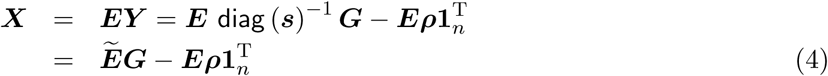

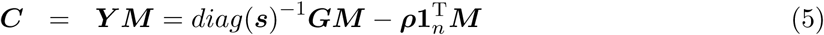

Here 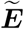 can be computed in time *𝒪*(*km*) while 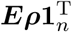 and 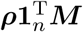 can be computed in time *𝒪*(*nk* +*mk*).

The key bottleneck in the E-step is the multiplication of the genotype matrix ***G*** by each of the *k* rows of the matrix 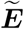 and in the M-step, multiplication of ***G*** by each of the *k* columns of the matrix ***M*** respectively. Leveraging the fact that each element of the genotype matrix ***G*** takes values in the set {0, 1, 2}, we can improve the complexity of these multiplication operations from *𝒪*(*nmk*) to 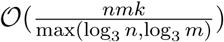 by extending the Mailman Algorithm [40]. For additional implementation details, see SI Section S1.

### The Mailman algorithm

In the M-step, we need to compute ***c*** = ***Ab*** for an arbitrary real-valued vector ***b*** and a *m* × *n* matrix ***A*** whose entries take values in {0, 1, 2}. We assume that *m* = ⌈log_3_(*n*)⌉. Naive matrix-vector multiplication takes 𝒪(⌈log_3_(*n*)⌉*n*) time.

The Mailman algorithm decomposes ***A*** as ***A*** = ***U***_*n*_***P***. Here ***U***_*n*_ is the *m* × *r* matrix whose columns containing all *r* = 3^*m*^ possible vectors over {0, 1, 2} of length *m*. We set an entry *P*_*i,j*_ to 1 if column *j* of ***A*** matches column *i* of 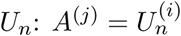. The decomposition of any matrix ***A*** into ***U*** _*n*_ and ***P*** can be done in 𝒪 (*nm*) time. Given this decomposition, the desired product ***c*** is computed in two steps, each of which has 𝒪 (*n*) time complexity [40]:

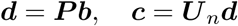

The Mailman algorithm provides computational savings in a setting where the cost of computing the decomposition of ***A*** are offset by the gains in repeated multiplication involving ***A***.

Similarly, in the E-step, we need to compute ***f*** ^T^***A*** in 𝒪(⌈log_3_(*n*)⌉*n*) time by computing ***A***^T^***f*** and computing a decomposition of ***A***^T^. A drawback of this approach is the need to store both decompositions that would double the memory requirements of the algorithm. Instead, we propose a novel variant of the Mailman algorithm that can compute ***f***^T^***A*** in 𝒪(⌈log_3_(*n*)⌉*n*) time using the same decomposition as ***A*** (SI Section S2).

Additional details on efficient implementation of the EM and Mailman algorithms can be found in SI Section S1.

### Simulations

We simulated genotypes at *m* independent SNPs across *n* individuals in which a single ancestral population diverged into *q* sub-populations with drift proportional to the *F*_*st*_, a measure of population differentiation. The allele frequency at SNP *f*_*j*,0_,*j* ∈ {1…, *m*} in the ancestral population was sampled from a uniform distribution such that 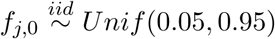. Allele frequencies in each of the *l* subpopulations were generated by simulating neutral drift from the ancestral allele frequency, 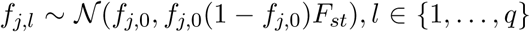 and were set to 0 or 1 if they fell outside the interval [0, 1]. The genotypes of an individual in population *l* at SNP *j* was sampled from a *Binomial*(2, *f*_*j,l*_) distribution.

### Benchmarking

To compare estimated PCs to reference PCs, we computed the mean of explained variance (MEV) – a measure of the overlap between the subspaces spanned by the two sets of PCs. Two different sets of *K* principal components each produce a K-dimensional column space. A metric for the performance of a PCA algorithm against some baseline is to see how much the column spaces overlap. This is done by projecting the eigenvectors of one subspace onto the other and finding the mean lengths of the projected eigenvectors. If we have a reference set of PCs (*v*_1_, *v*_2_, …, *v*_*k*_) against which we wish to evaluate the performance of a set of estimated PCs (*u*_1_, *u*_2_,…, *u*_*k*_), 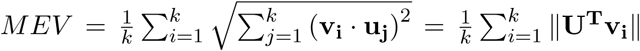 where U is a matrix whose column vectors are the PCs which we are testing.

In practice, when attempting to compute the top *k* PCs, ProPCA was found to converge faster by computing *l* PCs for *l*> *k* PCs and retaining the top *k* PCs. We set *l* = 2*k* in our experiments. While ProPCA could be run to convergence, we found that running it for *k* iterations already gave accurate results across the range of parameters considered. Our empirical results are consistent with our theoretical result that the EM algorithm converges exponentially fast in the spectral norm of the error matrix [31, 41] (SI Section S3).

We compared ProPCA to the current state-of-the-art methods for computing PCs from genotype data: the SVD implementation in PLINK (PLINK SVD [19]), FastPCA [12] and FlashPCA2 [13]. PLINK SVD refers to an exact computation of PCs using the full Singular Value Decomposition as implemented in the PLINK package (PLINK SVD). FastPCA [12] is an implementation of the randomized subspace iteration method [31] while FlashPCA2 [13] is an implementation of the implicitly restarted Arnoldi method [42]. We used default parameters for all methods. All experiments were performed on a Intel(R) Xeon(R) CPU 2.10GHz server with 128 GB RAM, restricted to a single core, capped to a maximum runtime of 100 hours and a maximum memory of 64 GB.

### Selection scan

The White British cohort was identified by the UK Biobank as participants who self-identified as ‘British’ within the broader-level group ‘White’ while having similar ancestral background [18]. For our selection scan, we further filtered the 409, 634 individuals in the White British subset to obtain an unrelated White British subset by removing individuals with one other related individual in the data set (individuals with kinship coefficients greater than 0.0625 (third-degree relatedness) to any other individual as determined by KING [43]). After removing these individuals, we obtained an unrelated White British subset containing 276, 736 individuals.

We inferred the top five PCs using ProPCA on all 276, 736 unrelated White British individuals and a filtered SNP set containing 146, 671 SNPs (UK Biobank SNP set). SNPs in the UK Biobank SNP set consist of SNPs on the UK Biobank Axiom array from which SNPs were removed if they have missing rates greater than 1.5%, minor allele frequencies (MAF) less than 1%, or if they were in regions of long-range linkage disequilibrium. The remaining SNPs were then pruned for pairwise *r*^2^ less than 0.1 using windows of 1000 base pairs (bp) and a step-size of 80 bp.

We developed a selection statistic to search for SNPs whose variation is not well-explained by the ProPCA model (closely related to the selection statistic proposed in [12]). Under the probabilistic PCA model, the normalized genotype matrix is modeled by a low rank approximation and Gaussian noise, **Y** = **CX** + ***ϵ***. Given our low rank approximation of the genotype matrix, **Ŷ** = **CX**, we have the residual: **Y** - **Ŷ** = ***ϵ***. For a SNP *j*, the Gaussian noise, ***ϵ***_***j***_ ∼ *𝒩*(**0**, *σ*^2^**I**_*n*_). Projecting this residual onto a PC results in a univariate Gaussian with zero mean and constant variance across SNPs. This variance can be estimated as the sample variance 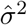 of the resulting statistics across SNPs. In summary, we propose the statistic: 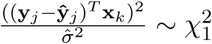 for SNP *j*, given the *k*-th PC. The projection of the residual onto a PC allows the signal of selection to be interpreted in the context of deviations from ancestry captured by the specific PC.

Furthermore, a variant of this statistic, which we call the combined statistic, can be generated from the selection statistics computed on each individual PC using the observation that the resulting chi-squared statistics are independent of each other. This allows us to create an additional statistic by summing the individual PC statistics to create a combined statistic that follows a chi-squared distribution with additional degrees of freedom for each PC used.

Using the results from the PCA on the UK Biobank SNP set, we performed our selection scan on a different set of 516, 140 SNPs. We generated this set of SNPs by removing SNPs that were multi-allelic, had genotyping rates less than 99%, had minor allele frequencies less than 1%, and were not in Hardy-Weinberg equilibrium (*p*< 10^*-*6^).

We performed an allele frequency test for each novel SNP using the Nomenclature of Territorial Units for Statistics level 3 (NUTS3) classification of regions for the UK. The NUTS3 classification defines non-overlapping borders for each region in the UK, allowing us to uniquely map each individual to a region in the UK using their birth location coordinates by checking which NUTS3 regions they fell into. For each of our novel loci, we then performed an two-tailed *Z*-test between each region’s allele frequency against all other regions. We corrected for multiple testing using the Bonferroni correction.

## Supplemental Data

Supplemental data includes 7 figures and 8 tables. 2 of 8 of the tables can be found in the accompanying Excel file.

## Acknowledgements

We would like to thank Bogdan Pasaniuc, members of his lab, and other members of the Sankararaman lab for advice and comments on this project. This research was conducted using the UK Biobank Resource under application 33127.

