## Supplementary Information for "Scalable probabilistic PCA for large-scale genetic variation data"

### Supplementary Information: Scalable and flexible Probabilistic PCA for large-scale genetic variation data

#### Contents


**S1 Implementation details** 1

**S2 Novel variant of the Mailman algorithm for left multiplication** 2

**S3 Convergence of ProPCA in the noiseless setting** 3

**S4 Exploring the contribution of the Mailman algorithm to scalability** 4

**S5 Application of ProPCA to missing data** 4

S5.1 PCA with Missing Data . . . . . . . . . . . . . . . . . . . . . . . . . . . . . 4

S5.2 EM for PCA with Missing Data . . . . . . . . . . . . . . . . . . . . . . . . . . 5

**S6 Application of ProPCA on 1000 Genomes Phase 1 Data** 5

**S7 White British Selection Scan and Analysis** 6


#### S1 Implementation details

**Application of the Mailman algorithm to the EM algorithm** For a genotype matrix $\boldsymbol{G}$ where $m > \lceil \log_3(n) \rceil$, we partition $\boldsymbol{G} = \left(\boldsymbol{G}_1^{\mathrm{T}} \ldots \boldsymbol{G}_B^{\mathrm{T}}\right)^{\mathrm{T}}$ into $B = \lceil \frac{m}{log_3(n)} \rceil$ sub-matrices each of size $\lceil log_3(n) \times n \rceil$ and decompose each $\boldsymbol{G}_b = \boldsymbol{U}_n \boldsymbol{P}_b$.

The M-step (Equation 5) requires computing $\boldsymbol{G\alpha}$ for $k$ distinct vectors $\boldsymbol{\alpha}$. We compute $\boldsymbol{G\alpha} = \begin{pmatrix} G_1\boldsymbol{\alpha} \\ G_2\boldsymbol{\alpha} \\ \vdots \\ G_B\boldsymbol{\alpha} \end{pmatrix}$. Since each of the products $\boldsymbol{G}_b\boldsymbol{\alpha}, b \in \{1, \ldots, B\}$ can be computed in $\mathcal{O}(n)$ operations (given $\boldsymbol{U}_n$, and $\boldsymbol{P}_b$), the entire matrix-vector product $\boldsymbol{G\alpha}$ can be computed in $\mathcal{O}(\frac{nm}{\log_3(n)})$ time.

The E-step (Equation 4) requires computing $\boldsymbol{\beta}^{\mathrm{T}}\boldsymbol{G}$ for $k$ distinct vectors $\boldsymbol{\beta}$. We compute this product as $\sum_{b=1}^{B} \boldsymbol{\beta}_b^{\mathrm{T}} \boldsymbol{G}_b$ in $\mathcal{O}(\frac{nm}{\log_3(n)})$ time where each term in the sum is computed using our novel variant of the Mailman algorithm.

**Likelihood Computation** To check for convergence, we need to compute the likelihood of the parameters in each iteration of the EM algorithm which is equivalent to the computing the squared Frobenius norm of the error matrix, *i.e.*, $||\mathbf{Y} - \mathbf{CX}||_F^2$.

$$
\begin{aligned}
||\mathbf{Y}-\mathbf{CX}||_F^2 &= tr[\ (\mathbf{Y}-\mathbf{CX})(\mathbf{Y}-\mathbf{CX})^T] \\
&= -2tr(\mathbf{Y^TCX}) + tr(\mathbf{X^TC^TCX}) + const
\end{aligned}
$$

Let $\boldsymbol{Z} = \boldsymbol{C}^{\mathrm{T}}\boldsymbol{Y}$. $\boldsymbol{Z}$ and $\boldsymbol{X}$ are $k \times n$ matrices so that the first term in the sum above ($tr(\boldsymbol{Z}^{\mathrm{T}}\boldsymbol{X})$) can be computed in $\mathcal{O}(nk)$ time. $\boldsymbol{Z}$ can be computed in $\mathcal{O}(\frac{nmk}{max(\log_3(n),\log_3(m)})$ using the Mailman algorithm. Thus, the likelihood can be computed in $\mathcal{O}(\frac{nmk}{max(\log_3(n),\log_3(m)} + nk)$.

We note that the columns of the Maximum Likelihood Estimate (MLE) of $\mathbf{C}$ do not correspond to the principal components of $\boldsymbol{Y}$ but instead span the principal subspace of the top $k$ eigenvectors of $\mathbf{Y}$. We can orthogonalize the matrix $\mathbf{C}$ to obtain the principal components in time $\mathcal{O}(mk^2)$, using *e.g.*, the Q-R decomposition.

**Efficient implementation of the Mailman algorithm** There are several considerations in an efficient practical implementation of the Mailman algorithm. While the multiplication with the $\boldsymbol{U}$ matrix is obtained by a recursion, we convert this into an iterative algorithm. Another important factor arises from the fact that the Mailman algorithm needs access to elements in the input vector that are not necessarily located in consecutive memory addresses. This can lead to frequent cache misses that can substantially reduce the efficiency of the implementation. To get around this limitation, we implemented a batched version of the Mailman algorithm. This version uses the idea that typically we need to multiply more than one vector at a time, *e.g.*, we often need to compute $k = 5$ PCs. Our implementation operates on the batch of input vectors at a time using the resulting locality among the input vectors. We use a default batch size of 10 although other batch sizes could also be used.

**Memory considerations** In the mailman algorithm, the matrix $\boldsymbol{U}_n$ is only used implicitly and need not be stored. The $\boldsymbol{P}$ matrix has the property that each column has exactly one entry that is one while all the other entries are zero. $\boldsymbol{P}$ can be stored as a length $n$ vector $\boldsymbol{p}$ indicating the locations of the one entry in each column of $\boldsymbol{P}$ . Since each element of the $\boldsymbol{p}$ vector is an integer, such that $p_i \in [1, n], i \in \{1, \ldots, n\}$, we can store $\boldsymbol{p}$ in $\lceil log_2(n) \rceil$ bits. This can be efficiently represented by storing 2 or more elements of $p$ in a single four byte integer. The storing and retrieval of an element can be performed by bit operations which increase the computational complexity moderately while reducing the memory requirements considerably.

#### S2 Novel variant of the Mailman algorithm for left multiplication

The EM algorithm requires alternate left and right multiplication of genotype matrix $\boldsymbol{G}$ in the E- and M-steps respectively. One approach to using the Mailman algorithm for each step consists of partitioning $\boldsymbol{G}$ along the columns and the rows respectively followed by computing decompositions of each of the resulting sub-matrices. This approach, however, doubles the memory requirement of the resulting algorithm. Instead, we propose a variant of the Mailman algorithm for left multiplication of a matrix with a vector that uses the same decomposition as for right multiplication.

Recall that for right multiplication, we would like to compute $\boldsymbol{c} = \boldsymbol{A}\boldsymbol{b}$ for an arbitrary real-valued vector $\boldsymbol{b}$ and a $m \times n$ matrix $\boldsymbol{A}$ whose entries take values in $\{0, 1, 2\}$. We assume that $m = \lceil \log_3(n) \rceil$. The Mailman algorithm decomposes $\boldsymbol{A}$ as $\boldsymbol{A} = \boldsymbol{U}_m\boldsymbol{P}$. Here $\boldsymbol{U}_m$ is the $m \times m_0$ matrix whose columns containing all $m_0 = 3^m$ possible vectors over $\{0, 1, 2\}$ of length $m$. $\boldsymbol{P}$ is a $m_0 \times n$ matrix. We set an entry $P_{i,j}$ to 1 if column $j$ of $\boldsymbol{A}$ matches column $i$ of $U_m$: $A^{(j)} = U_m^{(i)}$.

All other entries of $\boldsymbol{P}$ are set to zero. The decomposition of any matrix $\boldsymbol{A}$ into $\boldsymbol{U}_m$ and $\boldsymbol{P}$ can be done in $\mathcal{O}(nm)$ time. Given this decomposition, the desired product $\boldsymbol{c}$ is computed in two steps, each of which has $\mathcal{O}(n)$ time complexity [1]:

$$\boldsymbol{d} = \boldsymbol{P}\boldsymbol{b}, \quad \boldsymbol{c} = \boldsymbol{U}_m\boldsymbol{d}$$

We now describe an algorithm to compute $\boldsymbol{f}^{\mathrm{T}} = \boldsymbol{e}^{\mathrm{T}}\boldsymbol{A}$ using the same decomposition $\boldsymbol{A} = \boldsymbol{U}_m\boldsymbol{P}$. As in the setting of right multiplication, this algorithm proceeds in two steps:

$$\boldsymbol{g}^{\mathrm{T}} = \boldsymbol{e}^{\mathrm{T}}\boldsymbol{U}_m, \quad \boldsymbol{f}^{\mathrm{T}} = \boldsymbol{g}^{\mathrm{T}}\boldsymbol{P}$$

For the first step, we have:

$$\begin{aligned} \boldsymbol{g}^{\mathrm{T}} = \boldsymbol{e}^{\mathrm{T}}\boldsymbol{U}_m &= \begin{pmatrix} e_1 & \boldsymbol{e}_{2:m}^{\mathrm{T}} \end{pmatrix} \begin{pmatrix} \boldsymbol{0}_{3^{m-1}}^{\mathrm{T}} & \boldsymbol{1}_{3^{m-1}}^{\mathrm{T}} & \boldsymbol{2}_{3^{m-1}}^{\mathrm{T}} \\ \boldsymbol{U}_{m-1} & \boldsymbol{U}_{m-1} & \boldsymbol{U}_{m-1} \end{pmatrix} \\ &= \begin{pmatrix} \boldsymbol{e}_{2:m}^{\mathrm{T}}\boldsymbol{U}_{m-1} & e_1\boldsymbol{1}_{3^{m-1}} + \boldsymbol{e}_{2:m}^{\mathrm{T}}\boldsymbol{U}_{m-1} & e_1\boldsymbol{2}_{3^{m-1}}^{\mathrm{T}} + \boldsymbol{e}_{2:m}^{\mathrm{T}}\boldsymbol{U}_{m-1} \end{pmatrix} \end{aligned} \tag{1}$$

Here $e_1$ is the first element of $\boldsymbol{e}$ and $\boldsymbol{e}_{2:m}^{\mathrm{T}}$ is a vector of length $m-1$ consisting of elements 2 to $m$ of vector $\boldsymbol{e}$.

This gives us a recursive algorithm to compute $\boldsymbol{g}$ with base case :

$$\begin{aligned} e_m\boldsymbol{U}_1 &= e_m \begin{pmatrix} 0 & 1 & 2 \end{pmatrix} \\ &= \begin{pmatrix} 0 & e_m & 2e_m \end{pmatrix} \end{aligned} \tag{2}$$

The time complexity of this algorithm is given by $T(m) \leq 3^m + T(m-1) \leq 3^{m+1} = 3 \times 3^{\lceil \log_3(n) \rceil} = \mathcal{O}(n)$.

For the second step, note that each column of $\boldsymbol{P}$ has exactly one non-zero entry (with value equal to one). Thus, each entry of $\boldsymbol{f}$ can be computed in constant time so that $\boldsymbol{f}$ can be computed in $\mathcal{O}(3^m) = \mathcal{O}(n)$ time.

Thus, the total time complexity of computing $\boldsymbol{f}$ is $\mathcal{O}(n)$ instead of $\mathcal{O}(n\log_3(n))$ using naive matrix-vector multiplication.

For a general matrix $\boldsymbol{A}$ where $m > \lceil \log_3(n) \rceil$, we partition $\boldsymbol{A} = \begin{pmatrix} \boldsymbol{A}_1 \\ \boldsymbol{A}_2 \\ \vdots \\ \boldsymbol{A}_B \end{pmatrix}$ into $B = \lceil \frac{m}{log_3(n)} \rceil$ sub-matrices each of size $\lceil log_3(n) \times n \rceil$ and decompose each $\boldsymbol{A}_b = \boldsymbol{U}_n\boldsymbol{P}_b$. To now compute $\boldsymbol{f}^{\mathrm{T}} = \boldsymbol{e}^{\mathrm{T}}\boldsymbol{A}$, we compute $\sum_{b=1}^{B} \boldsymbol{e}_b^{\mathrm{T}}\boldsymbol{A}_b$. Each product can be computed in $\mathcal{O}(n)$ time so that $\boldsymbol{f}$ can be computed in $\mathcal{O}(\frac{nm}{\log_3(n)})$.

### S3 Convergence of ProPCA in the noiseless setting

There are several techniques to analyze the convergence properties of ProPCA. Under the assumption that the linear Gaussian model is true, convergence results of the EM algorithm can be invoked [2]. An alternate view of convergence in the setting where $\sigma^2 \to 0$ arises from viewing the EM updates as mathematically equivalent to alternating least squares [3]. In this view, we can show that the spectral norm of the reconstruction error, *i.e.*, the error between the data matrix $\boldsymbol{Y}$ and its rank-$k$ approximation $\boldsymbol{C}\boldsymbol{X}$, decreases to the optimal value at a rate that is exponential in the number of iterations. Our arguments follow from a combination of previous theoretical results.

The range of the matrix $\boldsymbol{C}^{(t)}$ obtained at the end of iteration $t$ of the EM algorithm is the same as the range of the matrix $\boldsymbol{Y}{\boldsymbol{Y}^{\mathrm{T}}}^{t}\boldsymbol{C}_0$ (Theorem 5 of Szlam et al. 2017). Setting $\boldsymbol{C}_0 = \boldsymbol{Y}\boldsymbol{\Omega}$ where $\boldsymbol{\Omega}$ where $\boldsymbol{\Omega}$ is a $n \times l$ matrix ($l = 2k$) with entries drawn independently from a standard normal distribution. Let $\boldsymbol{Q}^{(t)}$ denote the orthonormal basis for the range of $\boldsymbol{C}^{(t)}$. Then $\mathbb{E}\left[\|\boldsymbol{Y} - \boldsymbol{Q}^{(t)}{\boldsymbol{Q}^{(t)}}^{T}\boldsymbol{Y}\right]\| \leq (1+\alpha)^{\frac{1}{2t+1}}\sigma_{k+1}$ (Corollary 10.10 of Halko *et al.*, 2009). Here $\sigma_{k+1}$ is the $(k+1)^{st}$ largest singular value of $\boldsymbol{Y}$ and $\alpha$ is a constant that depends on the $m, n$ and $k$

### S4 Exploring the contribution of the Mailman algorithm to scalability

To explore the contribution of the Mailman algorithm to the scalability, we explored variants of the EM algorithm underlying ProPCA that differ in the implementation of the core genotype matrix-vector multiplication. In addition to the Mailman algorithm for genotype matrix-vector multiplication (EM-Mailman), we considered an implementation where the genotypes are stored as a matrix of doubles using the Eigen matrix library [4] (EM1) as well as another implementation where the genotypes are stored in a compact representation in which each genotype is represented using two bits (EM2). The representation in EM2 is expected to be memory-efficient relative to EM1. However, since EM1 represents genotypes directly as a matrix object in Eigen, we expect EM1 to be computationally more efficient. Figure S2 supports this intuition. EM1 could only be applied to sample sizes of up to $70,000$ before reaching our memory limit. While EM2 can run sample sizes up to $1,000,000$, it is more than two orders of magnitude slower than EM-Mailman. While EM1 is substantially faster, EM-Mailman is about three times faster. We expect that, even if memory were not a constraint, the Mailman algorithm would remain faster than the basic EM algorithm. We note that the Mailman algorithm is only 3-4 times faster than the basic EM algorithm instead of the log factor predicted by theory. We suspect that a reason for this gap is that the Mailman algorithm, as implemented, has not been optimized for specific computing architectures unlike standard matrix algorithms.

### S5 Application of ProPCA to missing data

#### S5.1 PCA with Missing Data

The use of a probabilistic model allows for handling missing entries in the genotype matrix. We assume that the genotype data is missing at random (MAR) [5], *i.e.*, the missingness depends only on the other observed values. We partition the observed data $\boldsymbol{G}$ into observed and unobserved entries. In the missing data setting, the observation model becomes:

$$\boldsymbol{g}_i|\boldsymbol{x}_i, \boldsymbol{\epsilon}_i \quad = \quad \boldsymbol{\mu} + \boldsymbol{C}\boldsymbol{x}_i + \boldsymbol{\epsilon}_i \tag{3}$$

Here $\boldsymbol{\mu}$ is a length $m$ vector denoting the mean genotype vector. Unlike the fully observed setting where the maximum likelihood estimate of $\boldsymbol{\mu}$ is equal to the sample mean $\overline{\boldsymbol{g}}$, in the missing data setting, we need to estimate $\boldsymbol{\mu}$ within the EM algorithm.

$$
\begin{aligned}
O &= \{\ (i,j)\ \mid\ \ g_{ij}\ is\ observed\ \},\\
O_j &= \{\ i\ \mid\ \ (i,j) \in O\ \},\\
O_i &= \{\ j\ \mid\ \ (i,j) \in O\ \},\\
\boldsymbol{x}_j &= j^{th}\ column\ of\ \mathbf{X},\\
\boldsymbol{c}_i &= i^{th}\ row\ of\ \mathbf{C}\ written\ as\ a\ column,\\
\mu_i &= the\ mean\ of\ g_{ij}\ where\ j \in O_i
\end{aligned}
$$

##### S5.2 EM for PCA with Missing Data

$$
\text{E Step:} \qquad \boldsymbol{x}_j = (\sum_{i \in O_j} \boldsymbol{c}_i \boldsymbol{c}_i^T)^{-1} \sum_{i \in O_j} \boldsymbol{c}_i (y_{ij} - \mu_i) \tag{4}
$$

$$
\text{M Step:} \qquad \boldsymbol{c}_i = (\sum_{j \in O_i} \boldsymbol{x}_j \boldsymbol{x}_j^T)^{-1} \sum_{j \in O_i} (y_{ij} - \mu_i) \boldsymbol{x}_j \tag{5}
$$

$$
\mu_i = \frac{1}{|O_i|} \sum_{j \in O_i} (g_{ij} - \boldsymbol{c}_i^T \boldsymbol{x}_j) \tag{6}
$$

Using the same ideas of the Mailman algorithm, the EM algorithm for missing data has a running time of $\mathcal{O}(\frac{nmk}{max(\log_3 n, \log_3 m)} + n_{missing} k^2)$ per iteration. Since the percentage of missing data is quite low, we can use the probabilistic model to efficiently handle missing data.

We evaluated the effectiveness of this extended model using simulated genotypes with missing data (Figure S3). We compared the accuracy of the PCs estimated using the extended model to the PCs estimated by running the EM algorithm on genotype data that was imputed through a random draw from a binomial distribution parameterized by the allele frequencies.

We simulated ten sets of complete genotypes with 50,000 SNPs and 10,000 individuals from 5 and 10 populations, each at differing $F_{ST}$ levels from 0.001 to 0.01 at intervals of 0.001. We simulated missing data by randomly removing 5% and 10% of the genotypes. To estimate the variance of our method, we averaged over 10 datasets.

For each method tested, we computed the MEV between the PCs inferred from the missing data and the PCs computed by applying SVD to the original genotype data with no missing values. Figure S3 shows that the PCs inferred from the ProPCA implementation that explicitly handles missing data are more accurate than the PCs computed by running ProPCA on imputed genotypes.

#### S6 Application of ProPCA on 1000 Genomes Phase 1 Data

We applied our method to genotype data from Phase 1 of the 1000 Genomes project [6]. On a dataset of $1,092$ individuals and $442,350$ SNPs, ProPCA computes the top five PCs with an MEV of 0.968 in about 17 seconds on a single core. The top two PCs computed by ProPCA and by running SVD on this data set are qualitatively indistinguishable (Fig. S4). On a dataset of $1,092$ individuals and about 33 million SNPs, ProPCA computes the top five PCs in 1240 seconds. Table S1 shows the runtimes of each of the methods tested (ProPCA, FastPCA, FlashPCA2 and PLINK_SVD) on the 1000 Genomes data and we observe that the ProPCA is computationally efficient.

### S7 White British Selection Scan and Analysis

Among the significant loci that we did not highlight in the main text, there are several genic loci have biological significance.

Transcriptome-wide association studies (TWAS) suggest that gene expression at *HERC6* is associated with gout ($p = 3.8 \times 10^{-123}$) [7]. Epidemiological studies in the UK also have shown that Wales, the geographic region associated with differences in *HERC6* allele frequencies, is among the regions of the UK with the highest prevalence and incidence in the UK [8]. The specific variant that is putatively under selection at this locus, rs112873858, does not appear to be significantly associated with gout in the UK Biobank however (logistic regression $p = 0.2395$).

*HERC2* (hect domain and RLD2), contains a single SNP in *HERC2* that is a primary determinant of light eye color in modern Europeans [9] and has been previously shown to be under selection [10]. A number of other SNPs in the *HERC2* locus have also been shown to be associated with iris color [11]. In the UK Biobank, we find that the SNP with the most significant p-value in *HERC2*, rs1129038, is associated with childhood sunburn occasions ($p = 6 \times 10^{-134}$) as well as skin and hair pigmentation ($p = 9.4 \times 10^{-103}$) (Table S6,S7).

*SKI* is a proto-oncogene located at a region close to the p73 tumor suppressor gene [12]. It is implicated in the TGF-$\beta$ signaling pathway [13] and has been shown to play a role in a variety of cancers [12, 14]. However, our specific locus does not appear to be significantly associated with any cancer in the UK Biobank.

Our combined selection statistic also resulted in an additional genic loci we did not highlight in the main text. The *AMPH* locus is located in the gene that codes for the amphiphysin protein, which is associated with the cytoplasmic surface of synaptic vesicles [15]. The gene is also implicated in stiff person syndrome and breast cancer [15]; however we were unable to find any significant associations with traits in the UK Biobank.

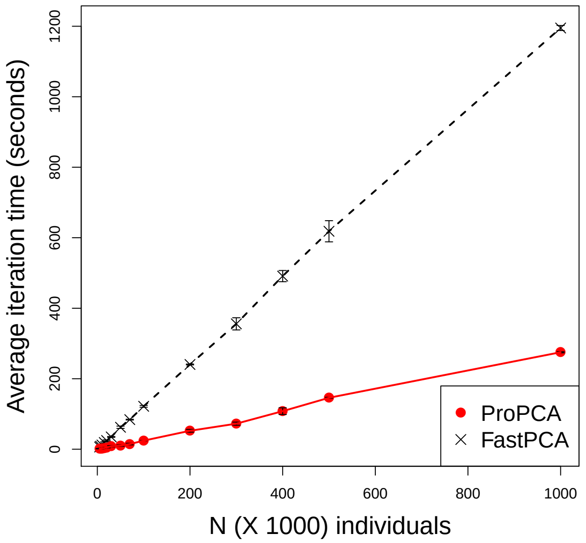


Figure S1: **ProPCA has faster per-iteration runtimes versus FastPCA**: Comparison of average per-iteration runtimes over simulated genotype data containing 100,000 SNPs, six subpopulations, $F_{st} = 0.10$ and individuals varying from 10,000 to 1,000,00. We were unable to leverage the source code for FlashPCA2 for this comparison.

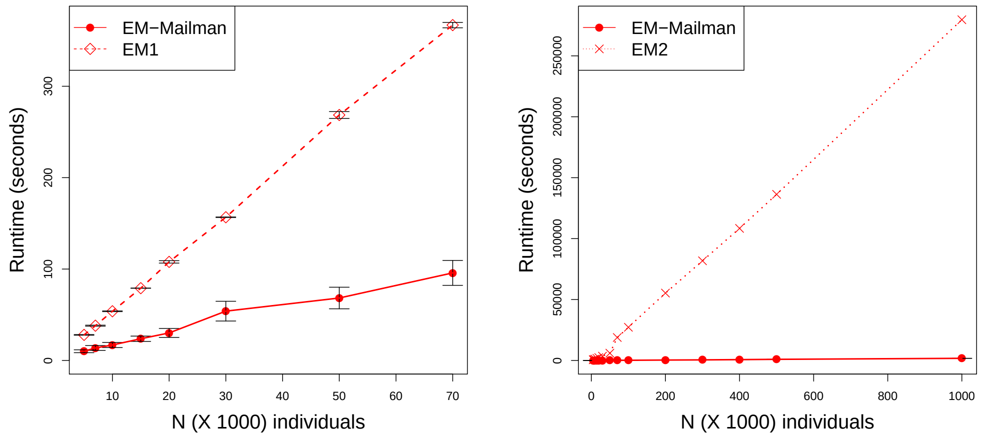


(a) (b)

Figure S2: **The Mailman matrix-vector multiplication contributes to the scalability of ProPCA**: We compare the time taken to compute the top five principal components by the EM algorithm underlying ProPCA when used in conjunction with the Mailman algorithm and without. We performed these comparisons on simulated genotype data containing 100,000 SNPs, six subpopulations, $F_{st} = 0.10$ and individuals varying from 10,000 to 1,000,000. Figures S2a compares the runtime of the EM algorithm with the Mailman matrix-vector multiplication to an EM algorithm where the genotypes are represented as a matrix of doubles (EM1). With this representation, the EM algorithm could only be applied to sample sizes of at most 70,000 individuals due to memory constraints. Figure S2b compares the runtime of the EM algorithm with the Mailman matrix-vector multiplication to an EM algorithm where genotypes are represented in a compact representation (EM2).

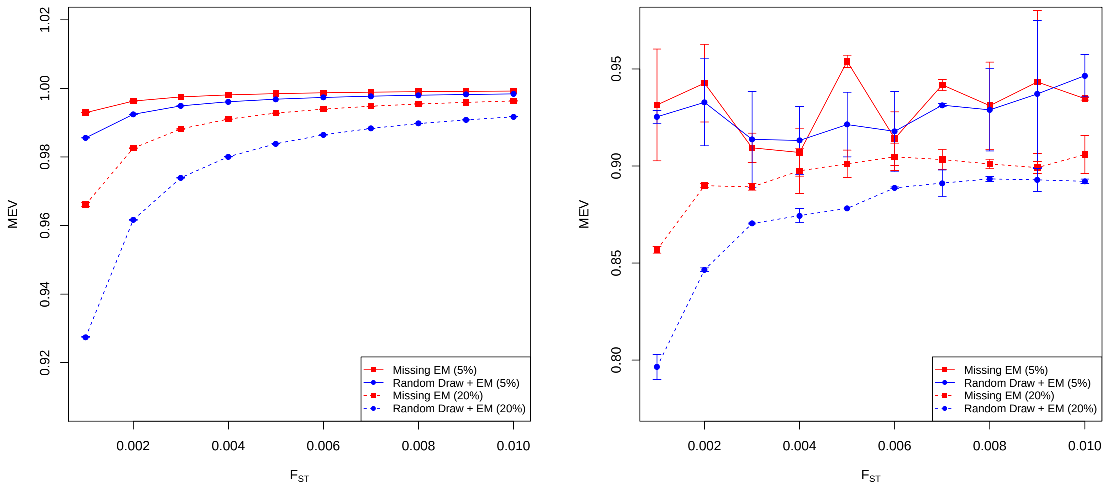


Figure S3: **ProPCA infers more accurate principal components (PCs) in the presence of missing data compared to imputed genotypes**: We compared the MEV from eigenvectors calculated from both modes of ProPCA with ground truth eigenvectors from performing a full SVD. We evaluated performance at 5% and 20% random missing values at 5 (left) and 10 principal components (right). The data consists of simulated genotype data of 50,000 SNPs from 10,000 individuals from 5 populations for 5 PCs and 10 populations for 10 PCs separated by a range of $F_{st}$ values. This process was repeated ten times to measure variability. Error bars denoting one standard deviation are shown for each point.

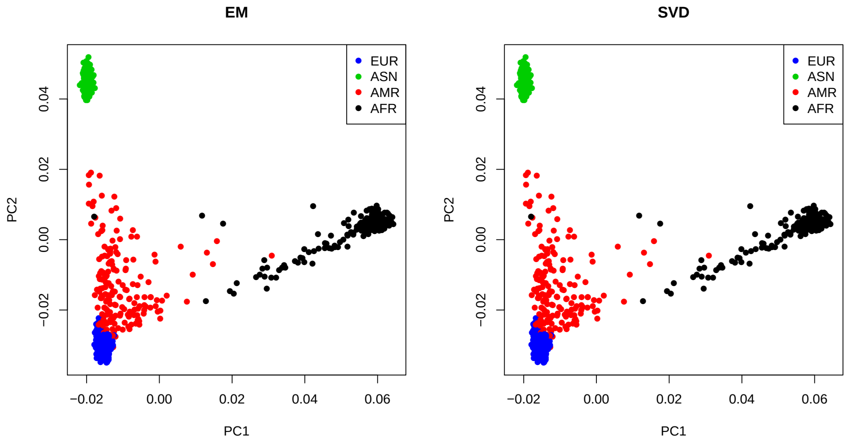


Figure S4: **ProPCA estimates principal components that are qualitatively indistinguishable from a full SVD on 1000 Genomes Phase 1 data.** EM refers to ProPCA.

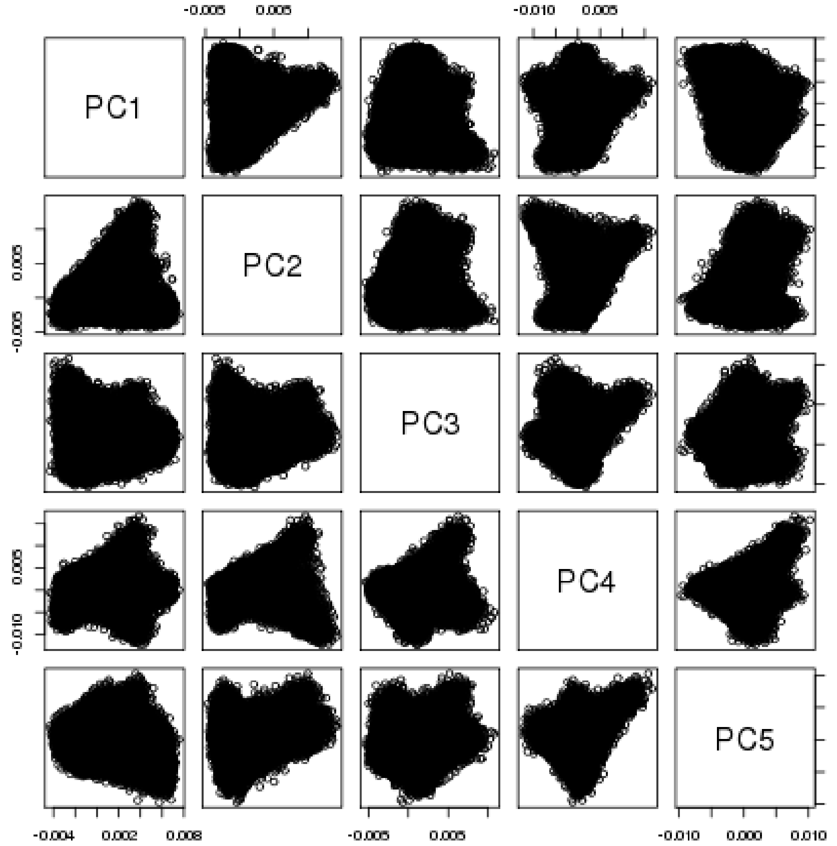


Figure S5: **Scatterplot pairs between the projections of the first five principal components of the unrelated White British**: Plotting pairs of the first five principal components reveals structure amongst the unrelated White British. This structure diminishes as we increase the number principal components used.

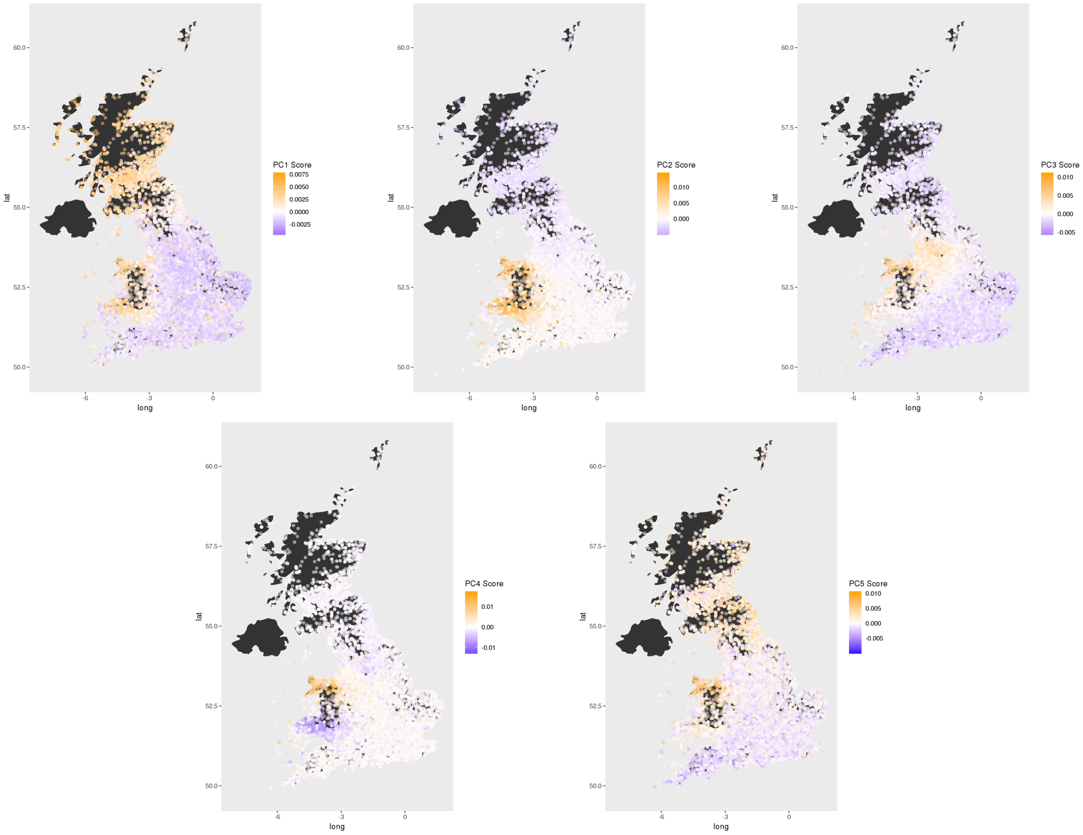


Figure S6: **Principal component scores of the unrelated White British overlaid on a map of the UK**: Using birth location data available in the UK Biobank, we placed a scatter plot colored by principal component score to reveal geographic variation captured by the principal components.

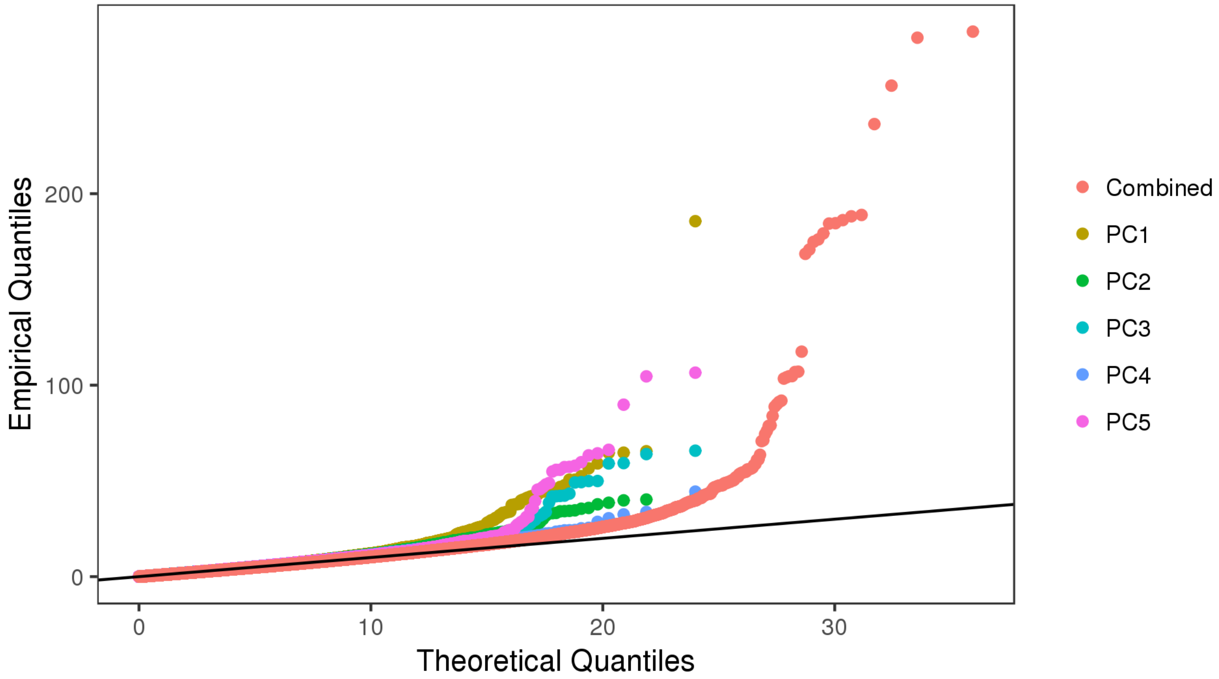


Figure S7: **The selection statistic is calibrated in the unrelated White British**: We plot the theoretical quantiles of the $\chi^2_1$ distribution against each of the empirical quantiles observed from the first five principal components. All five principal components follow the theoretical distribution well until the upper tail. We additionally show the calibration of the combined statistic against the theoretical quantiles of the $\chi^2_5$ distribution.

| Number of SNPs | Runtime (seconds) | | | |
|---|---|---|---|---|
| | ProPCA | FastPCA | FlashPCA2 | PLINK_SVD |
| $442,350$ | $\mathbf{17.1 \pm 0.9}$ | $49.0 \pm 1.5$ | $\mathbf{17 \pm 0.3}$ | 23.2 |
| $32,855,625$ | $\mathbf{1241.2 \pm 25.1}$ | NA | $3716.1 \pm 5.8$ | 1678.9 |

Table S1: **Comparison of runtime of methods to estimate principal components on the genotype data from the 1000 Genomes Phase 1 project**. We compared the runtime of the ProPCA algorithm, FastPCA, FlashPCA2 and PLINK_SVD when applied to 1092 individuals in the 1000 Genomes Phase 1 project. We report runtime in seconds averaged over ten trials and the standard deviation for each method (except PLINK_SVD which was run only once). FastPCA gave us a segmentation fault for the larger SNP set.

| | **PC1** | **PC2** | **PC3** | **PC4** | **PC5** |
|---|---|---|---|---|---|
| **Longitude** | -0.39 (0) | -0.06 (6.29e-188) | -0.18 (0) | 0.03 (4.88e-42) | -0.11 (0) |
| **Latitude** | 0.38 (0) | -0.41 (0) | 0.16 (0) | -0.05 (1.06e-141) | 0.32 (0) |

Table S2: **Pearson correlation between principal components and birth location coordinates in the unrelated White British.** Pearson correlation between the principal components and birth location coordinates reveals that the principal components unveil geographic variation. P-values from Pearson correlation $t$-test is shown in parentheses on the right.

| $\lambda_{GC}$ | **PC1** | **PC2** | **PC3** | **PC4** | **PC5** | **Combined** |
|---|---|---|---|---|---|---|
| **Statistic** | 0.961 | 0.970 | 0.979 | 0.955 | 0.958 | 0.962 |
| **Galinsky** | 1.017 | 0.904 | 0.900 | 0.791 | 0.794 | 0.877 |

Table S3: **Selection statistics are not substantially inflated.** We calculated the $\lambda_{GC}$ values for each principal component and the combined statistic to check for inflation. In the unrelated White British set, the calculated values show that our selection statistics are not substantially inflated (top row). Furthermore, we show that the previously related statistic proposed by Galinsky et al. 2016 does not calibrate as well as our statistics based on $\lambda_{GC}$ values (bottom row).

Table S4: **Table of significant SNPs found by selection scan on unrelated White British.** Our selection scan on the unrelated White British population resulted in 59 significant SNPs. Our significance threshold was a Bonferroni corrected $p < 0.05$. Bonferroni corrected $p$-values are shown.

| CHR | POS | rsid | Gene | PC1 | Other Gene Hits in Window |
|---|---|---|---|---|---|
| 2 | 135837906 | rs7570971 | RAB3GAP1 | 2.64E-09 | RAB3GAP1,R3HDM1,LCT |
| 4 | 38799710 | rs4833095 | TLR1 | 1.80E-09 | TLR10,TLR1,TLR6,FAM114A1 |
| 6 | 421281 | rs62389423 | | 8.80E-36 | IRF4,EXOC2 |
| 6 | 32139813 | rs9267817 | | 0.019121 | HLA |
| 19 | 49206417 | rs492602 | FUT2 | 0.002174 | FUT2 |
| **CHR** | **POS** | **rsid** | **Gene** | **PC2** | **Other Gene Hits in Window** |
| 1 | 2240074 | rs79907870 | SKI | 0.007996 | SKI |
| 1 | 116977051 | rs6670894 | | 0.015407 | |
| 4 | 38799710 | rs4833095 | TLR1 | 0.000341 | TLR10,TLR1,FAM114A1 |
| 5 | 164861910 | rs77635680 | | 0.000687 | |
| 9 | 13954710 | rs12380860 | | 0.026696 | |
| 15 | 28365618 | rs12913832 | HERC2 | 0.001613 | HERC2 |
| 16 | 53720436 | rs61747071 | RPGRIP1L | 0.006486 | RPGRIP1L |
| **CHR** | **POS** | **rsid** | **Gene** | **PC3** | **Other Gene Hits in Window** |
| 2 | 136407479 | rs1446585 | R3HDM1 | 0.001585 | R3HDM1,LCT |
| 4 | 38799710 | rs4833095 | TLR1 | 1.56E-09 | TLR10,TLR1,TLR6,FAM114A1 |
| **CHR** | **POS** | **rsid** | **Gene** | **PC4** | **Other Gene Hits in Window** |
| 4 | 89323743 | rs112873858 | HERC6 | 8.29E-05 | HERC6 |
| 5 | 164847509 | rs79194719 | | 0.019347 | |
| **CHR** | **POS** | **rsid** | **Gene** | **PC5** | **Other Gene Hits in Window** |
| 4 | 38798935 | rs5743614 | TLR1 | 1.75E-18 | TLR10,TLR1,TLR6,FAM114A1 |
| 6 | 421281 | rs62389423 | | 0.006962 | |

Table S5: **Principal component selection scan reveals 12 unique loci under selection across the top five principal components.** We obtained 59 selection hits across the first five principal components of the unrelated White British subset of the UK Biobank. We clustered these hits into 12 unique loci by aggregating all significant hits into 1 Mb windows centered around the most significant hits. Other genes with significant hits that are within the 1 Mb window are listed in the last column.

| SNP | Genes in Window | P | Phenotype |
|---|---|---|---|
| rs12913832 | HERC2 | 0 | pigment_HAIR_blackmale |
| | | 0 | pigment_HAIR_blonde |
| | | 0 | pigment_HAIR_darkbrown |
| | | 0 | pigment_HAIR |
| | | 9.70E-103 | pigment_HAIR_red |
| | | 0 | pigment_SKIN |
| | | 1.50E-138 | pigment_SUNBURN |
| | | 0 | pigment_TANNING |
| rs492602 | FUT2 | 2.50E-09 | blood_HIGH_LIGHT_SCATTER_RETICULOCYTE_COUNT |
| | | 1.80E-53 | blood_MEAN_PLATELET_VOL |
| | | 5.20E-11 | blood_MEAN_SPHERED_CELL_VOL |
| | | 9.70E-18 | blood_PLATELET_COUNT |
| | | 1.10E-08 | body_HEIGHTz |
| | | 7.50E-13 | bp_DIASTOLICadjMEDz |
| | | 1.20E-12 | bp_SYSTOLICadjMEDz |
| | | 9.40E-19 | disease_CARDIOVASCULAR |
| | | 2.60E-21 | disease_HI_CHOL_SELF_REP |
| | | 1.60E-09 | disease_HYPERTENSION_DIAGNOSED |
| | | 8.80E-12 | lung_FEV1FVCzSMOKE |
| rs62389423 | IRF4,EXOC2 | 6.80E-29 | blood_EOSINOPHIL_COUNT |
| | | 4.70E-19 | blood_LYMPHOCYTE_COUNT |
| | | 2.20E-16 | blood_WHITE_COUNT |
| | | 2.40E-68 | body_BALDING1 |
| | | 2.00E-66 | body_BALDING4 |
| | | 3.20E-31 | cancer_ALL |
| | | 0 | pigment_HAIR_blackmale |
| | | 0 | pigment_HAIR_blonde |
| | | 0 | pigment_HAIR_darkbrown |
| | | 0 | pigment_HAIR |
| | | 1.90E-33 | pigment_HAIR_red |
| | | 0 | pigment_SKIN |
| | | 0 | pigment_SUNBURN |
| | | 0 | pigment_TANNING |
| rs7570971 | RAB3GAP1,R3HDM1,LCT | 1.90E-08 | blood_EOSINOPHIL_COUNT |
| | | 1.70E-09 | blood_RED_COUNT |
| | | 2.60E-15 | lung_FVCzSMOKE |
| rs9267817 | HLA | 2.50E-10 | blood_MEAN_PLATELET_VOL |
| | | 6.30E-13 | blood_MONOCYTE_COUNT |
| | | 3.70E-16 | blood_RBC_DISTRIB_WIDTH |
| | | 1.00E-13 | body_HEIGHTz |
| | | 7.00E-10 | bp_SYSTOLICadjMEDz |
| | | 2.40E-13 | impedance_BASAL_METABOLIC_RATEz |
| | | 6.10E-27 | lung_FEV1FVCzSMOKE |

Table S6: **Selection hits are associated with phenotypes in the UK Biobank.** We ran genome-wide association tests for 64 phenotypes in the full release of the UK Biobank for each of our loci. Phenotypes shown reached a *p*-value of genome-wide significance level ($0.05 \times 10^{-6}$).

| SNP | Genes in Window | Phenotype Code | P | Phenotype |
|---|---|---|---|---|
| rs12913832 | HERC2 | INI1737 | 6.61E-24 | Childhood_sunburn_occasions |
| rs492602 | FUT2 | INI3064 | 1.28E-09 | Peak_expiratory_flow_(PEF) |
| | | INI50 | 2.00E-11 | Standing_height |
| | | HC269 | 1.55E-14 | high_cholesterol |
| | | HC273 | 4.35E-08 | essential_hypertension |
| | | HC357 | 3.96E-10 | duodenal_ulcer |
| | | INI1289 | 4.42E-09 | Cooked_vegetable_intake |
| | | INI20015 | 1.51E-08 | Sitting_height |
| | | INI24019 | 3.36E-09 | Particulate_matter_air_pollution_(pm10);_2007 |
| | | HC188 | 4.85E-17 | cholelithiasis/gall_stones |
| | | HC215 | 8.22E-13 | hypertension |
| | | HC225 | 6.19E-14 | cholecystitis |
| rs5743614 | TLR10,TLR1,TLR6,FAM114A1 | HC382 | 4.97E-11 | asthma |
| | | HC49 | 1.18E-14 | hayfever/allergic_rhinitis |
| | | INI24019 | 1.25E-64 | Particulate_matter_air_pollution_(pm10);_2007 |
| rs62389423 | IRF4,EXOC2 | INI30120 | 1.62E-11 | Lymphocyte_count |
| | | INI30150 | 1.17E-14 | Eosinophill_count |
| | | INI30210 | 4.74E-08 | Eosinophill_percentage |
| | | INI50 | 1.26E-08 | Standing_height |
| | | INI134 | 3.57E-09 | Number_of_self-reported_cancers |
| | | INI1737 | 8.62E-164 | Childhood_sunburn_occasions |
| | | INI1873 | 6.06E-18 | Number_of_full_brothers |
| | | INI24004 | 1.66E-12 | Nitrogen_oxides_air_pollution;_2010 |
| | | INI24006 | 7.33E-17 | Particulate_matter_air_pollution_(pm2.5);_2010 |
| | | INI24017 | 6.46E-13 | Nitrogen_dioxide_air_pollution;_2006 |
| | | cancer1003 | 2.41E-87 | skin_cancer |
| | | cancer1060 | 2.05E-99 | non-melanoma_skin_cancer |
| | | FH1001 | 3.98E-18 | Lung_cancer |
| rs7570971 | RAB3GAP1,R3HDM1,LCT | INI3062 | 1.11E-08 | Forced_vital_capacity_(FVC) |
| | | INI23100 | 1.37E-08 | Whole_body_fat_mass |
| | | INI24019 | 5.36E-12 | Particulate_matter_air_pollution_(pm10);_2007 |
| rs9267817 | HLA | INI30100 | 1.36E-10 | Mean_platelet_(thrombocyte)_volume |
| | | INI30150 | 3.16E-16 | Eosinophill_count |
| | | INI46 | 1.31E-09 | Hand_grip_strength_(left) |
| | | INI50 | 1.51E-24 | Standing_height |
| | | HC303 | 5.00E-49 | malabsorption/coeliac_disease |
| | | INI20015 | 1.63E-14 | Sitting_height |
| | | INI21002 | 7.44E-12 | Weight |
| | | INI23098 | 2.31E-11 | Weight |
| | | INI24019 | 1.33E-10 | Particulate_matter_air_pollution_(pm10);_2007 |
| | | FH1065 | 7.46E-11 | High_blood_pressure |
| | | HC215 | 3.20E-10 | hypertension |

Table S7: **Selection hits are associated with phenotypes from the Global Biobank Engine.** We queried the Global Biobank Engine for associations from our loci. The Global Biobank Engine contains GWAS results for many more phenotypes than those available in the UK Biobank. Phenotypes shown are significant at genome-wide significance level ($0.05 \times 10^{-6}$).

| CHR | POS | rsid | Gene | P | Other Gene Hits in Window | PC1 | PC2 | PC3 | PC4 | PC5 |
|---|---|---|---|---|---|---|---|---|---|---|
| 1 | 2240074 | rs79907870 | SKI | 0.046830 | SKI | 1.27 | 35.48 | 4.63 | 2.15 | 1.38 |
| 2 | 136407479 | rs1446585 | R3HDM1 | 1.66E-14 | RAB3GAP1,R3HDM1,UBXN4,LCT | 56.55 | 2.40 | 38.63 | 2.12 | 5.02 |
| 3 | 54077256 | rs9856661* | | 0.020002 | | 0.94 | 9.18 | 12.43 | 0.72 | 23.45 |
| 4 | 89323743 | rs112873858 | HERC6 | 2.18E-05 | HERC6 | 0.02 | 7.65 | 2.70 | 44.40 | 6.36 |
| 4 | 38799710 | rs4833095 | TLR1 | 6.22E-53 | TLR10,TLR1,TLR6,FAM114A1 | 65.50 | 41.64 | 65.78 | 7.13 | 104.60 |
| 5 | 162948205 | rs116352364* | | 0.000162 | | 0.00 | 13.68 | 21.83 | 16.15 | 5.27 |
| 5 | 164861910 | rs77635680 | | 3.81E-11 | | 2.81 | 40.26 | 5.00 | 32.52 | 8.16 |
| 6 | 32526736 | rs111586361 | HLA-DRB5 | 0.003826 | HLA-DRB5 | 24.43 | 0.29 | 11.32 | 1.25 | 12.94 |
| 6 | 421281 | rs62389423 | | 7.11E-47 | IRF4,EXOC2 | 185.64 | 13.97 | 12.09 | 8.97 | 35.75 |
| 7 | 38463542 | rs118079376* | AMPH | 0.000817 | AMPH | 1.08 | 0.01 | 6.36 | 18.80 | 27.26 |
| 9 | 120475302 | rs4986790* | TLR4 | 2.35E-05 | TLR4 | 20.37 | 25.98 | 1.68 | 0.67 | 12.29 |
| 15 | 28365618 | rs12913832 | HERC2 | 0.012037 | HERC2 | 0.07 | 38.60 | 5.57 | 3.42 | 0.16 |
| 16 | 53720436 | rs61747071 | RPGRIP1L | 0.012396 | RPGRIP1L | 0.76 | 35.88 | 4.67 | 6.39 | 0.05 |
| 19 | 49206417 | rs492602 | FUT2 | 0.000505 | FUT2 | 38.02 | 12.08 | 0.33 | 3.03 | 1.08 |

Table S8: **Combined selection statistic across the top five principal components reveals four additional novel loci.** We discover four additional novel loci using our combined selection statistic from the first five principal components. Loci not found in the individual PC selection statistics are denoted by an asterik in the rsid column. The chi-squared statistic (one degree of freedom) for each principal component is shown in the last five columns of the table.

Table S9: **Allele frequency tests between NUTS3 regions at novel loci confirms differences between geographic regions.** We performed a two-tailed proportion test for our novel loci between the allele frequency in each individual region from the NUTS3 classification of the United Kingdom against the frequency from every other region. We corrected the $p$-values using the Bonferroni correction (11 loci $\times$ 163 regions). The corrected $p$-values for regions passing the significance threshold are shown in the table.
